# Branch-specific plasticity explains distal enrichment of retinotopically displaced inputs in visual cortex

**DOI:** 10.64898/2026.04.01.715858

**Authors:** Andrew T. Landau, Bernardo L. Sabatini, Claudia Clopath

**Author notes:** Correspondence |.

## Abstract

Neurons distribute synaptic inputs across their dendritic tree. In layer 2/3 pyramidal cells of primary visual cortex, spines on distal dendrites share somatic orientation preference but have receptive fields displaced in retinotopic space, which supports tuning to visual edges. However, it is not known how synaptic plasticity rules can lead to specialization of tuning properties across dendritic compartments. We demonstrate an experimentally grounded model of compartment-specific spike-timing dependent plasticity (STDP) that accounts for the enrichment of retinotopically-displaced inputs on distal branches. Our previous experimental work revealed compartment-specific calcium signals that predict reduced STDP-mediated depression but preserved potentiation. Based on these findings, we built an STDP model with compartment-specific properties, in which some distal branches are relatively resistant to STDP-mediated depression. Synapses on these branches are more likely to stabilize inputs with weaker correlations to postsynaptic spiking. Using a visual input model, we show that compartment-specific reduction in STDP-mediated depression recapitulates in vivo experimental measurements of spine tuning. Furthermore, our experimental results show that reduced STDP-mediated depression is restricted to distal dendritic compartments with complex branching structure and not observed in other distal branches. Therefore, our model makes an untested prediction that complex branches will be hotspots for retinotopically-displaced inputs.

## Introduction

Neurons segregate synaptic inputs in different compartments of their dendritic tree. For example, bottomup sensory inputs tend to arrive on proximal sites near the soma, whereas top-down contextual inputs tend to arrive on distal sites within apical branches [1; 2; 3; 4]. This organization is determined by an interplay of anatomy and synaptic plasticity. Anatomy constrains where synaptic connections can be made, and plasticity sculpts the strength and selectivity of potential connections.

Compartment specific plasticity rules support the functional segregation of inputs [5; 6; 7; 8; 9]. Although variation in plasticity rules is partially due to molecular differences across compartments, some variation is inherited from the electrical properties of the dendritic tree. In particular, the attenuation of back-propagating action potentials (bAPs) in distal compartments can shift the dynamics of global plasticity signals involved in spike-timing dependent plasticity (STDP) [10]. This can lead to variation in the timing rules of STDP [10] and engage local process that shift the balance from depression to potentiation [11].

Studies of primary visual cortex have identified that retinotopically displaced inputs, which are central to the formation of visual edge tuning, preferentially form on distal dendritic branches of layer 2/3 pyramidal cells [2; 12]. Computational studies have identified normative rules that produce visual edge tuning, ranging from simple source separation models like independent component analysis to more sophisticated models of efficient coding [13; 14]. However, these models fail to account for the dendritic distribution of inputs, instead representing neurons as single point compartments.

It is not known if the distribution of synaptic inputs in primary visual cortex can be explained by compartment-specific plasticity rules. Most studies of compartment-specific plasticity have focused on abstract functional benefits rather than specific input patterns [15; 16; 17]. Studies that aim to explain compartment specific input organization tend to focus on major differences between anatomical input classes, like comparing the subcellular targets of thalamic sensory inputs and long-range feedback connections [3; 18; 19; 20]. However, in the case of visual edge tuning, plasticity rules appear to be fully responsible for segregating input types in distinct dendritic compartments.

We propose an experimentally motivated and biophysically grounded model of compartment specific STDP that leads to a specific prediction about the distribution of synaptic inputs in layer 2/3 visual cortex cells and can explain existing measurements. In previous experimental work, we observed that sections of the dendritic tree with complex branching structure have a selective reduction in bAP-evoked calcium influx, which is preserved in distance matched branches with simpler morphology [21]. However, bAP-evoked amplification of synaptic calcium influx through NMDA-type glutamate receptors (NMDARs) is preserved in both branches.

In this paper, we demonstrate how our experimental measurements predict branch-specific variation in the ratio of STDP potentiation and depression (D/P ratio) in the distal dendritic tree [22]. Next, we show that the ratio of potentiation and depression shapes how correlated input synapses must be with each other to maintain stable connections [23]. We then demonstrate how this property directly produces the observed distribution of synaptic tuning in primary visual cortex [2]. Specifically, the high depression in proximal sites leads to highly correlated, retinotopically superimposed inputs, whereas the reduced depression in distal sites permits weakly correlated inputs that share the proximal orientation tuning but are retinotopically displaced. Our model can explain the observed distribution of spine tuning in visual cortex and makes an additional prediction that retinotopically displaced input are preferentially found on distal branches with complex branch structure.

## Results

### Branch-specific calcium signaling

Our previous experimental study uncovered branch-specific calcium signaling known to drive activity-dependent synaptic plasticity [21; 22]. We measured calcium influx in cortical layer 2/3 cells evoked by single back-propagating action potentials (bAPs), synaptic input, and synaptic input paired with a bAP delayed by 5ms (Figure 1a). We stimulated precisely timed bAPs with somatic current injection through a whole-cell patch pipette and stimulated synaptic input in individual dendritic spines with two-photon glutamate uncaging. In distal apical branches (>100*µ*m), we observed large variation in bAP-evoked calcium influx (Δ[*Ca*]_*AP*_, Figure 1a-c), despite controlling for distance from the soma. Regardless of Δ[*Ca*]_*AP*_ amplitude, all branches displayed large bAP-evoked amplification of synaptic calcium influx (Δ[*Ca*]_*amp*_, Figure 1a-d). Because we cannot directly control the concentration of glutamate released by uncaging, we plot relative Δ[*Ca*]_*amp*_, computed as the ratio of Δ[*Ca*]_*amp*_ and Δ[*Ca*]_*glu*_. This ratio measures how much bAPs amplify calcium influx through voltage-dependent NMDA receptors (NMDARs) due to transient relief of the *Mg*^2+^ block.

**Figure 1.**
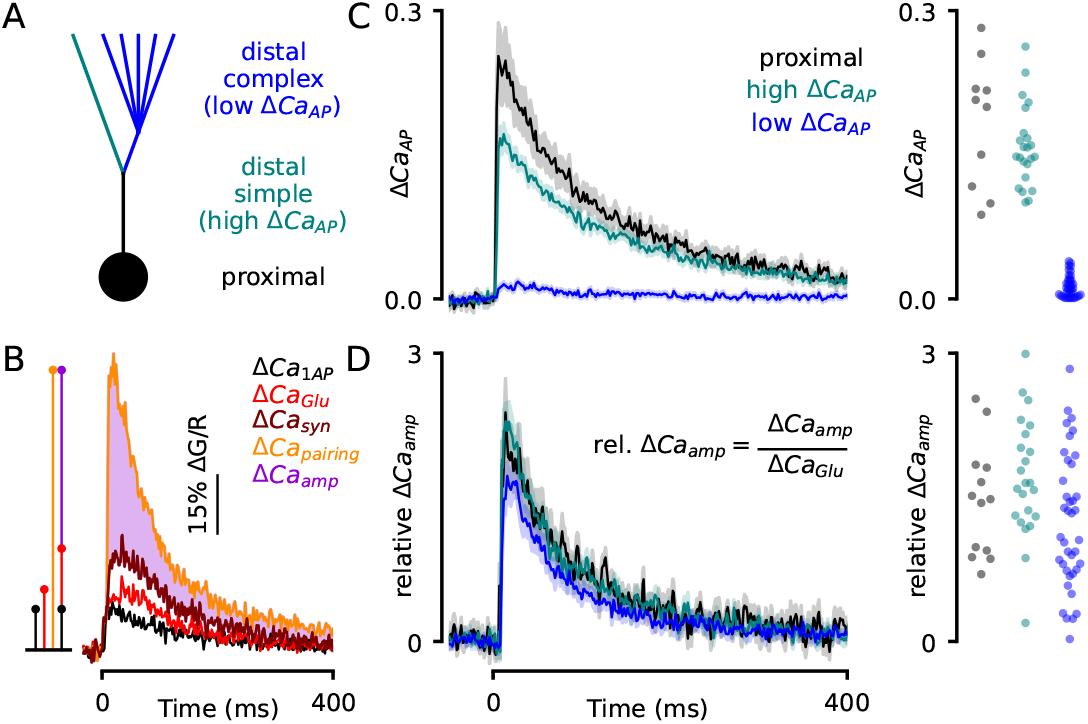
Mismatch in bAP-evoked calcium influx across dendritic branches. **(A)** Schematic of naming scheme for dendritic branches. Distal-complex and distal-simple are defined based on the amplitude of Δ[*Ca*]_*AP*_. **(B)** Example of evoked calcium signals. Δ[*Ca*]_*amp*_ is measured as the difference between the calcium evoked by pairing a bAP with glutamate uncaging and the synthetic linear sum of each independently. **(C)** Left: average Δ[*Ca*]_*AP*_ traces in each compartment (compartments defined by Δ[*Ca*]_*AP*_ amplitude). Right: peak amplitude across dendritic branches. **(D)** Left: average relative Δ[*Ca*]_*amp*_ in each compartment. Relative signal measured as Δ[*Ca*]_*amp*_*/*Δ[*Ca*]_*glu*_ to account for variation in NMDAR activation. Right: peak amplitude across branches.

Using dendritic loose-patch recordings, voltage imaging, and biophysical modeling, we deduced that variation in dendritic branch structure determines the amplitude of Δ[*Ca*]_*AP*_ (Figure 1a). Sections of the dendritic tree with simple branch structure have higher impedance, leading to high amplitude local bAPs that evoke calcium influx. Sections with more elaborate branch structure have lower impedance, which reduces the amplitude of local bAPs, so they fail to evoke calcium influx. Based on this morphological distinction, we refer to distal recording sites with high Δ[*Ca*]_*AP*_ as “distal-simple” and sites with low Δ[*Ca*]_*AP*_ as “distal-complex” (Figure 1a, and Figs 4-7 of Ref. [21]).

The mismatch in Δ[*Ca*]_*AP*_ and Δ[*Ca*]_*amp*_ observed in distal-simple and distal-complex sites predicts a divergence in the amount of depression and potentiation evoked by STDP protocols in each branch type. The magnitude of depression evoked by STDP is proportional to the amplitude of calcium influx mediated by voltage-gated calcium channels (VGCCs; Figure 2e,f) [22], which we measure as Δ[*Ca*]_*AP*_ . The magnitude of potentiation evoked by STDP is proportional to the amplitude of calcium influx mediated by NMDA-type glutamate receptors (NMDARs; Figure 2e,f) [22], which we measure as relative Δ[*Ca*]_*amp*_. Therefore, we hypothesize that distal-simple branches have typical levels of depression and potentiation, whereas distal-complex branches have reduced depression but typical potentiation.

**Figure 2.**
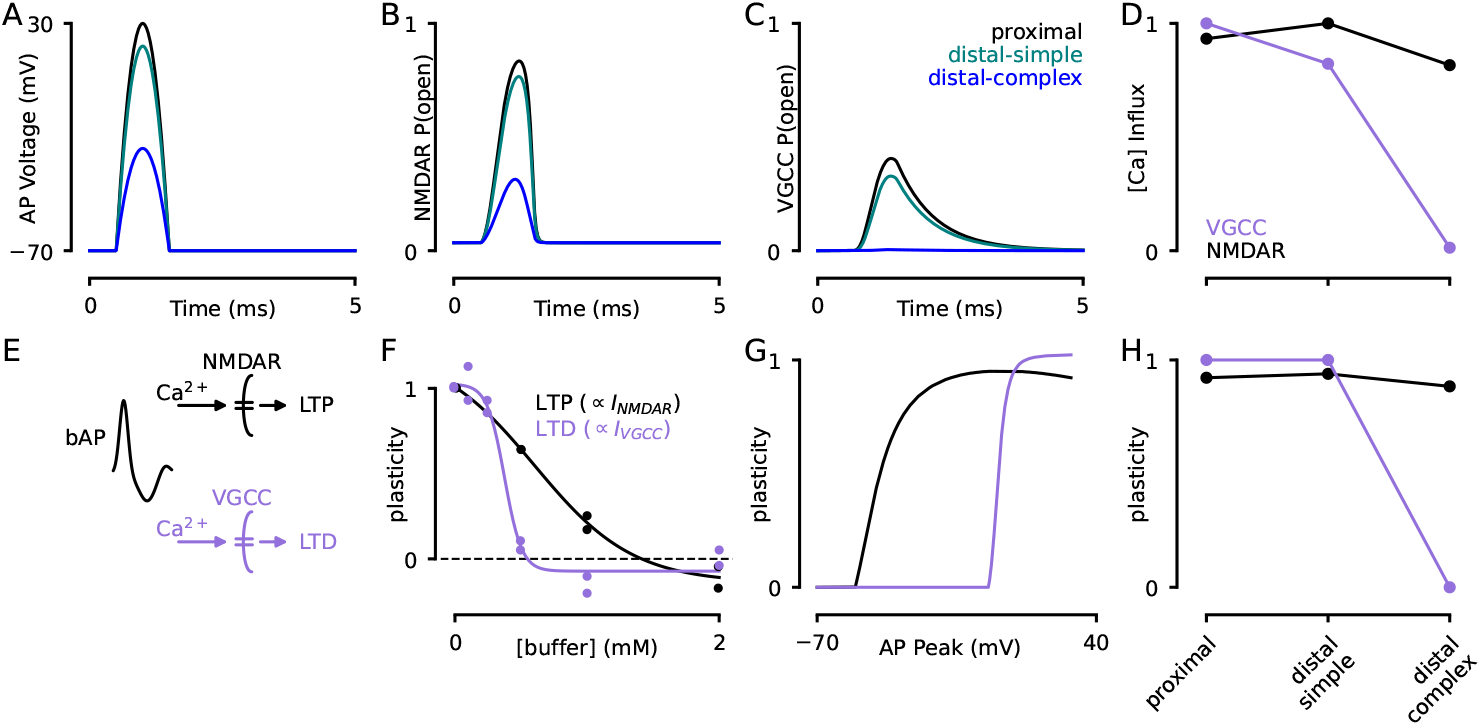
STDP Depression and Potentiation evoked in each branch type. **(A)** Action potentials were modeled as quadratic waveforms with variable peak amplitude to model experimental observations in the three dendritic branch sites. **(B)** NMDAR open-probability traces evoked by bAPs in each branch-type. **(C)** VGCC open-probability traces evoked by bAPs in each branch-type. **(D)** Integrated calcium influx in NMDARs and VGCCs evoked in each branch type. We used the GHK current equation to estimate dynamic calcium influx. **(E)** In STDP protocols, calcium influx through NMDARs and VGCCs leads to potentiation and depression, respectively. **(F)** Results from Nevian & Sakmann (2006, reproduced from published figure via plot digitization), showing how the magnitude of potentiation and depression depend on the amplitude of calcium influx. **(G)** Estimated potentiation and depression evoked by STDP as a function of local bAP amplitude. **(H)** Predicted magnitude of potentiation and depression evoked in each branch type, using the calcium influx measured in simulations from panels A-D.

To illustrate how differences in local bAP amplitude affect calcium influx through VGCCs and NMDARs, we simulated calcium currents evoked by bAPs using a biophysical model based on published work [24; 25]. Based on our loose-patch dendritic recordings and voltage-imaging data [21], we used bAPs with three different amplitudes to model distinct dendritic sites: 100 mV bAPs modeled proximal sites, 90 mV bAPs modeled distal-simple branches, and 45 mV bAPs modeled distal-complex branches (Figure 2a). Because of the shallow voltage dependence of NMDARs, lower amplitude bAPs were still able to open NMDARs in distal-complex branches (Figure 2b). However, because of the sharp, high-threshold voltage dependence of VGCCs, lower amplitude bAPs completely failed to open VGCCs in distal-complex branches (Figure 2c). We used the Goldman-Hodgkin-Katz current equation to measure the integrated calcium current evoked in each of these sites (Figure 2d). These results show why there is a mismatch in Δ[*Ca*]_*AP*_ and Δ[*Ca*]_*amp*_ in distal-complex branches, but not in distal-simple branches.

We built a simple model to predict the magnitude of depression and potentiation evoked by bAPs of different amplitudes. First, we built a transfer function that relates the magnitude of calcium influx through each channel type (VGCCs & NMDARs) to the magnitude of plasticity evoked by STDP. In a previous study [22], Nevian & Sakmann measured how much the magnitude of STDP decreases in the presence of calcium buffers that linearly reduce calcium concentration (Figure 2d). Because the free calcium concentrations that cause plasticity are so much lower than the buffer concentration, we used the buffer capacity formula to convert the concentration of buffer to relative calcium concentration [26], with a value of 1.0 representing the concentration without any calcium buffer. Next, we related the calcium evoked by different bAP amplitudes in our simulation (Figure 2d) to the relative calcium concentration in the transfer function. Because our simulation measures current per conductance density, and the conductance density of VGCCs and NMDARs are unknown, we scaled our data by the maximum calcium signal evoked by VGCCs or NMDARs experimentally [21]. As a result, we inferred the relationship between the peak bAP amplitude and the potentiation or depression evoked by STDP (Figure 2f,g). Finally, we show the resulting magnitude of potentiation and depression evoked in proximal, distal-simple, or distal-complex sites (Figure 2h). These simulations confirm our hypothesis that the mismatch in calcium signaling maps to a divergence in plasticity in each branch type.

To explore how the divergence in depression and potentiation affects the tuning properties of synapses in distal-simple or distal-complex branch types, we built a model of STDP that incorporates the properties of each branch type. We used an additive model of plasticity from Song & Abbott [23] that increments the strength of synapses based on the timing of preand post-synaptic spikes (Figure 3a). This model requires the strength of depression to exceed potentiation to prevent runaway firing. Because we wanted to explore how reducing depression affects function, we included a homeostatic scaling rule, which multiplicatively scales all synapses whenever a neuron is above or below its firing set point (Figure 3b) [27].

**Figure 3.**
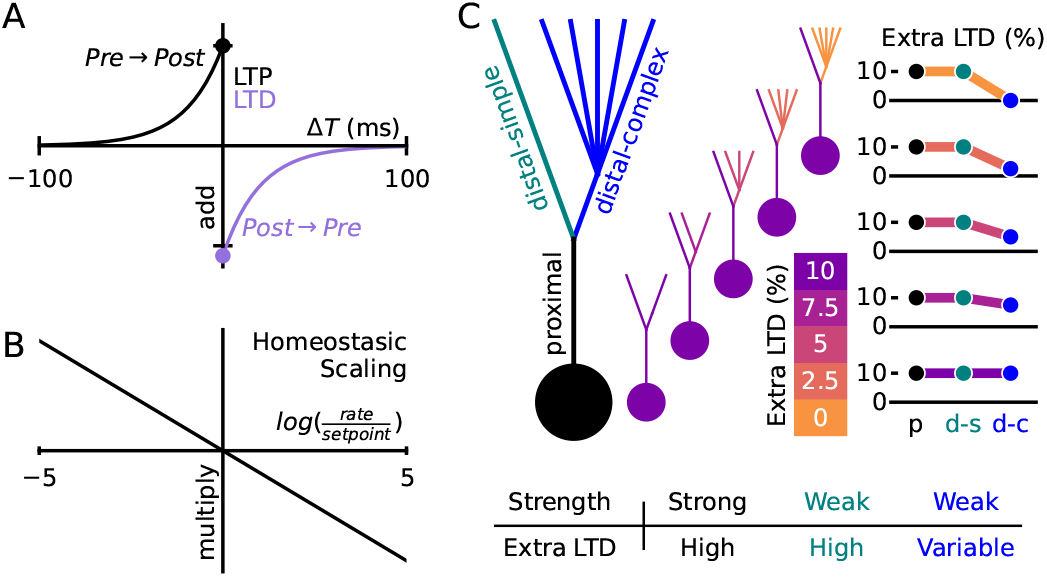
Branch-specific STDP Model. **(A)** Additive STDP plasticity rule. Pre→post pairings evoked potentiation, and post→pre pairings evoked depression. **(B)** Homeostatic plasticity rule. Neurons adjusted all synapses multiplicatively in proportion to the log of the ratio between their current firing rate and their set point. **(C)** Branch-specific plasticity scheme. Each neuron we simulated had a proximal, distal-simple, and distal-complex compartment. The maximum allowed strength of synapses was high in proximal sites and low in both distal sites. We varied the amount of extra synaptic depression (LTD) in distal-complex branches.

Inspired by the results from Figure 2, we designed a three-compartment neuron in which each compartment varies in both the strength of synapses due to the distance from the soma [6] and the magnitude of excess depression due to the dendritic branch structure [21] (Figure 3c). Proximal sites had strong synapses and high extra depression. Distal-simple sites had weak synapses and high extra depression. Distal-complex sites had weak synapses and variable levels of extra depression, which represents the primary experimental axis that we explore.

Because STDP promotes correlated input tuning [23], we hypothesized that a reduction in synaptic depression would stabilize synapses with weaker correlations to other inputs. To test this hypothesis, we trained neurons on a simulated dataset in which each input synapse’s firing rate was partially correlated with a single latent source (Figure 4a,b), ranging from correlation values of 0 to 0.4. The correlation coefficient matrix of the input rates is shown in Figure 4b. We simulated STDP until neurons reached steady state, then analyzed the resulting synaptic weight on each input as a function of the dendritic compartment and the input correlation. Proximal sites developed strong tuning to highly correlated inputs, similar to ref [23] (Figure 4c,d, top). Distal-simple sites were weakly tuned to all inputs, because their weaker synaptic strength prevents them from reliably spiking the postsynaptic neuron (Figure 4c,d, middle). Distal-complex sites developed strong tuning to correlated inputs; however, as we varied the level of extra depression in distal-complex sites from 10% to 0%, the tuning curve shifted such that inputs with weaker correlations developed strong tuning (Figure 4c,d, bottom). We determined the half-point of sigmoidal fits to each tuning curve to quantify how correlated each input must be to acquire stable tuning (Figure 4e, middle inset). In proximal and distal-simple sites, the value was high and independent of the level of extra depression in distal-complex sites (Figure 4e, top & middle). In distal-complex sites, reducing extra depression led to a strong reduction in the half-point, such that the postsynaptic cell was tuned to inputs with random activity at 0% extra depression (Figure 4e, bottom).

**Figure 4.**
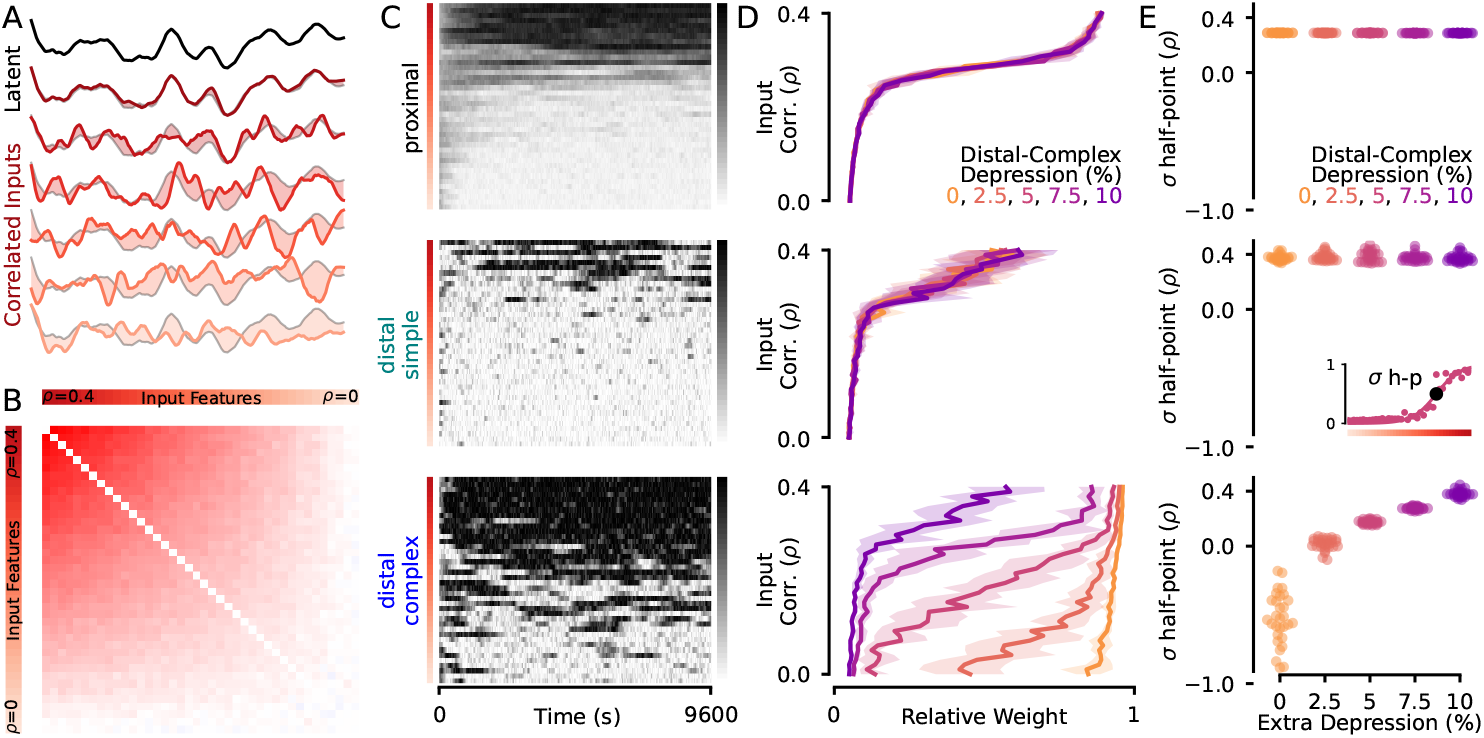
Strength of depression shapes input correlation properties. **(A)** Illustration of latent source model. A single autocorrelated latent source was generated. The dynamics of each input was partially determined by the latent source and partially by autocorrelated independent noise. The signal of each input is shown in color, and the difference from the latent source is shown by shading. **(B)** Resulting correlation matrix of all inputs (N=40) with correlation ranging from 0 to 0.4. **(C)** Total synaptic strength on each input in proximal (top), distal-simple (middle), and distal-complex (bottom) sites from a single recording. Each row represents the timecourse of synaptic weight on each presynaptic input source. **(D)** Average asymptotic synaptic strength as a function of input correlation. Weights were normalized to be relative to the maximum possible value. Color-coding follows Figure 3c, the color indicates the extra LTD magnitude in the distal-complex branch within each neuron. **(E)** Correlation of input source required to maintain a synaptic weight at 50% of the maximum value for each branch type, denoted “*σ* h-p”. Middle inset: how the *σ* h-p was measured. The level of extra depression in distal-complex sites had a large effect on *σ* h-p locally, but did not affect proximal or distal-simple sites.

Based on these simulations, we hypothesize that our compartment-specific STDP model can explain the distribution of input tuning in cortical layer 2/3 cells of mouse primary visual cortex [2]. To explore this possibility, we built a toy model of visual inputs to cortex in which a single postsynaptic cell receives input from a 3×3 grid of orientation tuned presynaptic cells (Figure 5). In each visual frame, the pixels could take on one of 4 orientations (Figure 5a). We presented a visual “edge” with probability *p*(*edge*), in which case an edge was formed by three pixels with matching orientations, always passing through the central pixel (Figure 5a). Otherwise, all pixels were random and independent. Presynaptic input cells had pre-determined tuning and were localized to single pixels, with von-mises tuning for the orientation in their pixel (Figure 5b). In the postsynaptic cell, proximal sites were connected to the central pixel, to model precise thalamocortical input as observed experimentally [20], while distal sites were connected to all pixels (Figure 5c). We used a weighted sum of Gabors to visually represent the tuning of each neuron, which showed that postsynaptic neurons became tuned to a single orientation that was coherent across the proximal, distalsimple, and distal-complex sites (Figure 6a,b).

**Figure 5.**
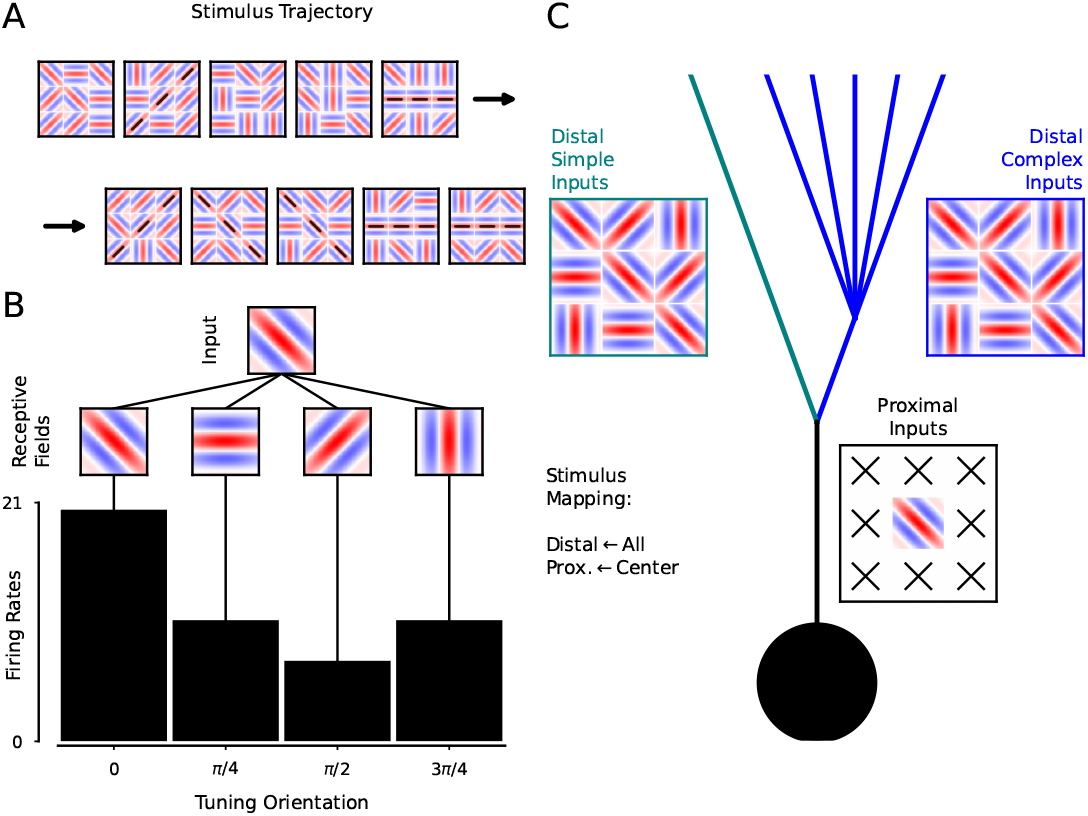
Visual edge tuning model. **(A)** The visual environment was a 3×3 grid of 4 possible orientations. Edges appeared randomly with probability *p*(*edge*) that were centered on the central pixel (black lines indicate edge). Pixels not participating in an edge had random orientations. **(B)** Presynaptic inputs followed von-mises tuning for orientation. The firing rate of each presynaptic input was entirely determined by a single pixel, so there were 36 total presynaptic input classes. **(C)** Compartment-specific input mapping. Proximal synapses were only connected to the 4 central pixel presynaptic inputs. Distal synapses were connected to all 36 presynaptic inputs.

**Figure 6.**
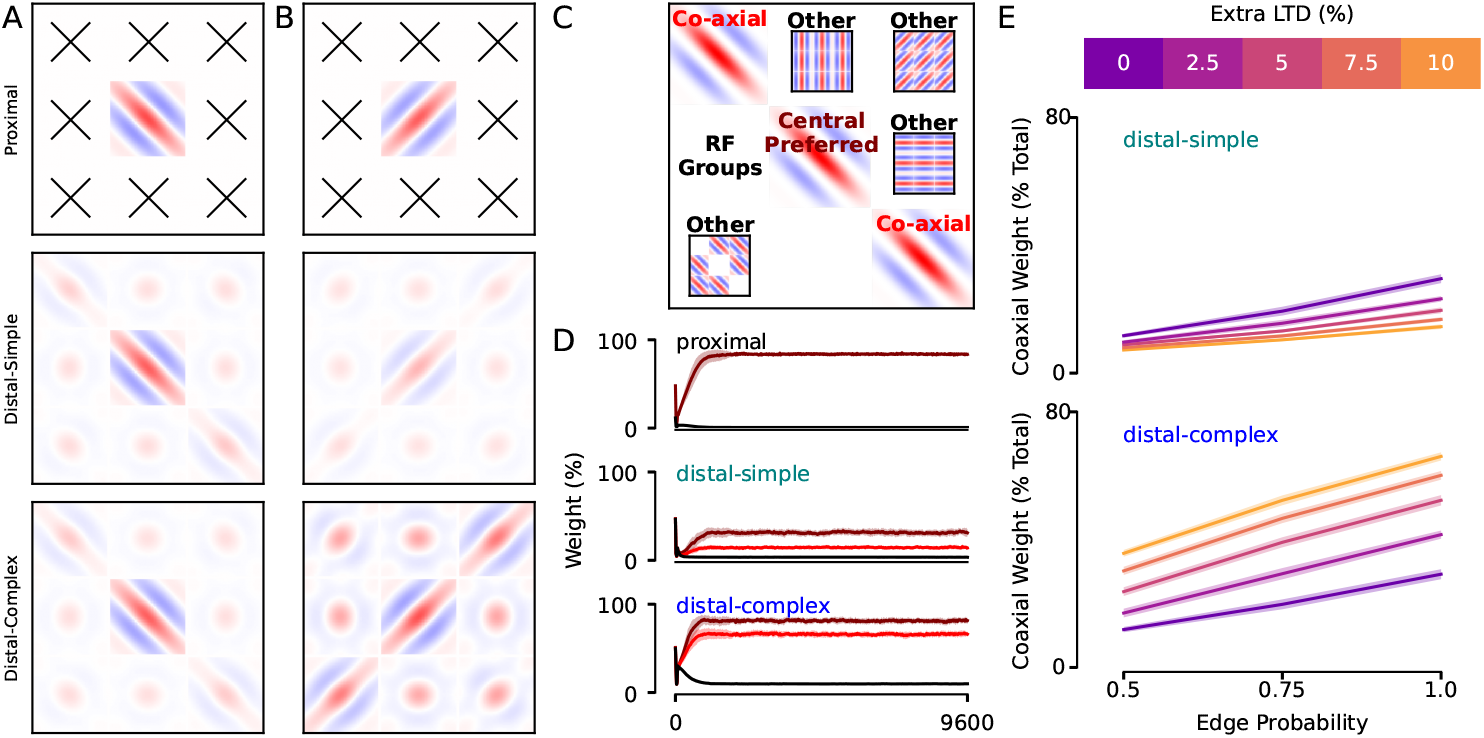
Recapitulation of experimental measurements of spine tuning. **(A)** Example of tuning at the end of simulation for a neuron with **high extra LTD** in the distal-complex branch. Both distal sites matched proximal tuning, with a hint of edge tuning. We used a weighted-sum of Gabors to represent tuning visually. **(B)** Example of tuning at the end of simulation for a neuron with **low extra LTD** in the distal-complex branch. The distal-complex branch shows a pronounced and selective increase in edge tuning. **(C)** Schematic of each receptive field group. Central-preferred refers to the orientation in the central pixel that matched proximal tuning. Co-axial tuning refers to the two boundary pixels that complete the edge of the central-preferred input. Other refers to all other presynaptic inputs (average of 33 other input types). **(D)** Compartment-specific tuning to each RF group across a full simulation. Y-axis shows magnitude of tuning relative to maximum possible value for each group. The simulation had *p*(*edge*)=1.0 and 0% extra LTD in the distal-complex site. **(E)** Average coaxial weight percentage across all simulations (N=30 each) as a function of *p*(*edge*) and extra LTD % in the distal-complex site.

In Iacaruso et al. [2], distal dendritic sites were more likely to have spines that shared orientation preference with the soma, but with retinotopically-displaced receptive fields. In our model, we observed this phenomenon in distal-complex sites depending on the level of extra depression and the edge probability. When extra depression was high, the distal-complex compartment had weak tuning to the retinotopically-displaced inputs (Figure 6a). However, when extra depression was low, it became strongly tuned such that its inputs represent the full edge (Figure 6b). To analyze this systematically, we divided inputs into three receptive field groups. “Central-preferred” inputs are in the central pixel and the preferred orientation of the postsynaptic cell (Figure 6c). “Coaxial” inputs are in the two outer positions that complete the edge corresponding to the central tuning type (Figure 6c). These are the key input types to analyze. “Other” inputs are all other inputs to the cell (Figure 6c).

All three compartments became strongly tuned to the central-preferred input (Figure 6d,e). However, as predicted by our hypothesis, the postsynaptic cells only acquired consistent tuning to coaxial inputs in the distal-complex site with reduced synaptic depression (Figure 6d,e). Coaxial tuning was more pronounced when visual edges were more frequent and depended on a reduction in synaptic depression (Figure 6d,e). Interestingly, the coaxial tuning of the distal-simple branch was suppressed by a reduction in extra LTD in the distal-complex branch due to the extra synaptic drive and homeostatic compensation (Figure 6e). Together, these results demonstrate that our compartment-specific STDP model can explain why retinotopically displaced (“coaxial”) inputs are preferentially localized on distal dendritic branches. Weak correlations are stable on distal branches in which the pressure to maintain high correlation from bAP-evoked synaptic depression is selectively reduced. Our model predicts that these inputs will not be on every distal branch but rather will specifically be in sections of the distal dendritic tree with complex branch structure, which causes the reduction in synaptic depression.

## Discussion

In this work, we provide a mechanistic explanation for how synaptic plasticity rules result in retinotopically displaced inputs becoming preferentially localized on distal dendritic branches of cortical layer 2/3 pyramidal cells of visual cortex [2]. Our model proposes causal links from dendritic biophysics to functional tuning properties by identifying how branch-specific calcium signaling affects STDP rules. Reductions in synaptic depression caused by attenuated local bAP amplitude allow weakly correlated inputs to maintain stable tuning [21]. Due to the correlation structure of visual input, this permits distal branches to become tuned to displaced inputs that play a central role in creating tuning to visual edges.

Our model recapitulates the distribution of dendritic spine tuning in layer 2/3 cortical pyramidal neurons of visual cortex. Furthermore, it makes a specific, untested prediction about the branch-specific distribution of displaced tuning in layer 2/3 cortical pyramidal neurons. Our experimental work shows that only sections of the dendritic tree with complex branch structure have a reduction in bAP-evoked calcium influx, which is required for reducing synaptic depression and stabilizing retinotopically displaced inputs [21]. Therefore, we predict that analysis of the morphological structure of layer 2/3 apical branches will reveal that displaced inputs preferentially appear on apical sites with complex branching.

Although our model was inspired by the observed distribution of compartment-specific spine tuning in visual cortex, it makes predictions that extend beyond that context. Our results with an abstract latent variable model in Figure 4 shows that reductions in synaptic depression promote weakly correlated synapses in general. While this applies to edge tuning in visual cortex, it may explain other tuning features in visual cortex or different sensory areas. Weakly correlated synapses are a fundamental component to multimodal tuning, in which the synapses across input modes are not as correlated with each other as synapses within an input mode. Vanilla STDP would predict that one input mode would eventually “win out” [23]. Our model provides an explanation for how multiple input modes can be preserved within a single cell. Furthermore, it predicts that dendritic spines tuned to a secondary input mode, like movement tuning in primary visual cortex, are preferentially observed on branches with reduced synaptic depression [19; 28]. As above, this is testable by comparing spine tuning properties with local dendritic branch structure.

We prioritized a simple model that is sufficient to explain the observed distribution of spine tuning in visual cortex. However, our model can be extended to include more complex or biophysically realistic plasticity rules. For example, voltage-dependent STDP may specifically interact with morphology of dendrites beyond the effect on AP-evoked calcium influx we modeled here [29]. Additionally, our model ignores local plasticity mechanisms like dendritic spikes that can produce even more compartment-specific clustering of synaptic input tuning, which may be exaggerated in distal-complex branches [30].

Together, our modelling shows that compartment-specific plasticity rules determined by the biophysical properties of dendrites can explain the observed distribution of synaptic tuning in visual cortex. Our model makes specific predictions about the branch-specific distribution of retinotopically displaced inputs that can be tested in future experiments. More broadly, our work shows how considering the physical structure of neurons can lead to insights about the computational properties of dendrites.

## Methods

### Experimental Data, Figure 1

Experimental data was collected as part of a previous study [21]. We refer the reader to that paper for a detailed description of the experimental methods. Here, we provide a brief summary of the methods relevant to the data in Figure 1. Data was collected from acute coronal slices of C57Bl/6j wild-type mice between postnatal days P21-P28. Somatic whole-cell recordings were acquired from cortical layer 2/3 pyramidal cells to measure intracellular voltage and elicit APs through brief current injection. We measured calcium influx through two-photon imaging of Fluo-5F and stimulated synaptic input through two-photon glutamate uncaging. In pairing protocols, APs were evoked 5ms after glutamate uncaging. Synthetic linear sums were computed by adding the baseline-subtracted calcium signal evoked by APs or glutamate uncaging alone, with a 5ms offset to reflect the timing of pairing protocols.

### Simulations, Figure 2

We simulated calcium influx through VGCCs and NMDARs using published data on the voltage-dependent open probability and time constants of each channel type [24; 25]. For VGCCs, we used the following voltage-dependent forward and backward rate constants of the activation gate (*α*_*m*_ and *β*_*m*_) and inactivation gate (*α*_*h*_ and *β*_*h*_):

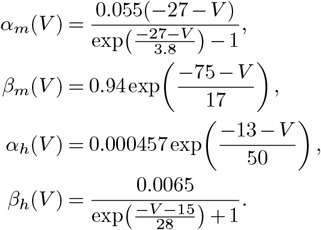

To compute open probability and time constant for the activation (denoted *m*) and inactivation (denoted *h*) gates of VGCCs, we used:

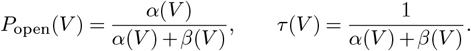

For NMDARs (denoted *n*), we calculated voltage-dependent open probability and time constant as:

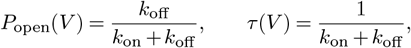

based on the measured on/off rates of the Mg^2+^ block [24], with [Mg^2+^] = 1 *m*M:

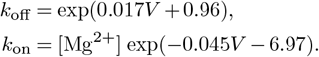

We used Euler’s method to compute changes in the state of each gate in dynamic simulations:

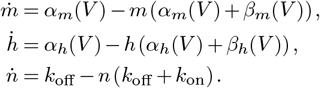

The open probability of NMDARs is equal to *n*, and the open probability of VGCCs is equal to *m*^2^*h*. To convert open probability into a voltage-dependent calcium current, we used a modified Goldman-Hodgkin-Katz current equation with [Ca]_in_ = 75 nM and [Ca]_out_ = 1.5 mM:

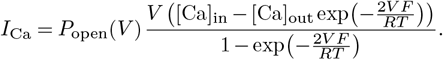

We used quadratic voltage depolarizations to mimic the shape of APs. For each stimulus, we used a preset amplitude (*V*_amp_) and duration (*V*_dur_), and solved for *a*:

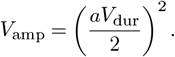

This gives a waveform over the domain −*V*_dur_*/*2 *< t < V*_dur_*/*2:

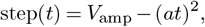

which we added to a baseline voltage of −70 mV. We used a 100mV AP in proximal compartments, a 90mV AP for distal-simple compartments, and a 45mV AP for distal-complex compartments. Although this was not measured directly in experiments, it matches the relative AP amplitudes measured with voltage imaging (Figure 5 of [21]) and in our biophysical simulations (Figure 7 of [21]).

To determine the integrated calcium influx in Figure 2d, we integrated the calcium current evoked by the voltage step after subtracting the baseline current (NMDARs are partially open at rest).

We used plot digitization to extract the data from Figure 4 of [22] and replotted it in Figure 2f.

To generate the transfer functions in Figure 2g, h, we first simulated AP-evoked calcium influx across a range of AP amplitudes (0–100 mV from a −70 mV baseline) and measured integrated calcium for NMDAR and VGCC currents separately. The Nevian/Sakmann curve in Figure 2f gives plasticity as a function of added buffer concentration, not directly as a function of calcium. To convert this into a transfer function of the form *Plasticity* = *f* ([*Ca*]_*relative*_), we used mass action to model relative calcium as a function of added buffer. At steady-state, the relationship between buffer concentration and free calcium concentration follows: 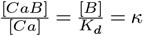. Buffer dynamics are fast compared to calcium influx and extrusion, so we can assume steadystate. Additionally, we assume that [*B*] doesn’t appreciably change due to calcium binding (which is true given the small changes in calcium concentration [26]). Therefore, we can solve for the fraction of free calcium remaining as a function of added buffer:

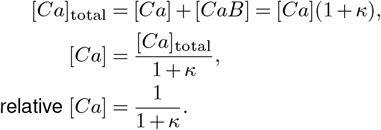

Our estimates of total integrated-calcium estimates for NMDAR and VGCC are proportional to the peak conductances of each channel type, which is not known. However, using the relative calcium equation as above, we can scale the curves to be relative to the maximum calcium influx observed through NMDARs and VGCCs. To do this, we estimated the relative peak calcium influx through NMDARs and VGCCs by comparing the maximal nonlinear calcium component and maximal AP-only calcium signal in the eLife dataset (approximating maximal NMDAR- and VGCC-mediated calcium influx, respectively), and used this ratio to scale the NMDAR-driven curve so both channel transfer functions reflected the same relative calcium influx.

### Branch specific STDP model, Figures 3–6

We modeled each neuron as a leaky integrate-and-fire unit receiving excitatory and inhibitory conductance-based synaptic input, following the model in [23]. The membrane voltage evolved as

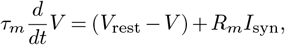

with spike generation at a fixed threshold and reset to *V*_rest_ after each spike.

Synaptic current was computed as

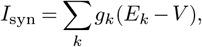

where *g*_*k*_ is synaptic conductance and *E*_*k*_ is the reversal potential of synapse type *k*. Conductance decayed exponentially and increased on presynaptic spikes in proportion to the weight of the synapse:

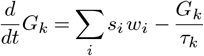

where *s*_*i*_(*t*) ∈ {0, 1} is a presynaptic spike and *w*_*i*_ is synaptic weight. Any synapse whos synaptic weight was below a fixed threshold *w*_*threshold*_ did not contribute to synaptic current

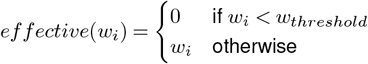

We used additive STDP with separate potentiation and depression eligibility traces. *A*_+_ and *A*_*−*_ are the potentiation and depression increments, respectively, which represent the maximum change in synaptic weight by a STDP pairing. Let

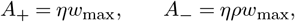

where *η* is the STDP rate and *ρ* is the depressionto-potentiation ratio. *w*_max_ is the maximum synaptic weight, such that additive increments to the weight were proportional to the range of possible synaptic weights. In general, *ρ* must be greater than 1 to ensure stability of the neuron [23].

STDP is implemented in three update rules. (1) Each neuron maintains a global depression trace that increments upon each postsynaptic spike (*s*_*post*_) and has exponential decay with time constant *τ*_*−*_. (2) Each synapse maintains a potentiation trace that increments upon each presynaptic spike and has exponential decay with time constant *τ*_+_. (3) Each synapse’s weight is updated by the depression trace whenever a presynaptic spike occurs, and by the potentiation trace whenever a postsynaptic spike occurs. Weights are clipped to [0, *w*_max_].

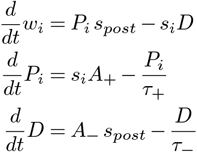

To stabilize firing rates, we included a multiplicative homeostatic plasticity rule, inspired by [27]. Neurons had a set point firing rate, *r*_set_ and a dynamic firing rate estimate 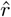 that evolved slowly over time (much slower than the time step and STDP timescales) with exponential decay and incremented by the instantaneous firing rate (*r*_*inst*_, where Δ*t* is the time step). The homeostatic drive was computed as the log ratio of the rate estimate and the set point, and synapses were multiplicatively updated by the homeostatic drive. Both equations use *τ*_*h*_ which is much slower than all other time constants used in the model (20 seconds).

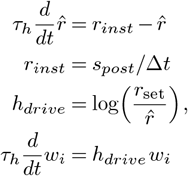

We modeled three excitatory compartments: proximal, distal-simple, and distal-complex following the three branch types identified in our experimental data [21]. All used the same STDP and homeostatic update equations, but differed in maximum synaptic weight (*w*_max_) and depression-to-potentiation ratio (*ρ*). Proximal synapses were stronger, reflecting their proximity to the soma. Distal-simple and distal-complex had equivalent reductions in max synaptic weight. Proximal and distal-simple synapses had higher depression-to-potentiation ratios, and therefore stronger LTD. We varied the depression-to-potentiation ratio (referred to as “Extra LTD%”) specifically in distal-complex synapses to create a branch-specific condition of extra LTD.

### Latent variable STDP environment, Figure 4

To determine how branch-specific depression shapes the input correlation properties of a neuron learning with STDP, we simulated neurons in a latent-variable environment in which the presynaptic inputs had a rank 1 correlation structure along with isotropic noise. We generated a single, autocorrelated latent source. Then, the dynamics of each presynaptic firing rate was partially determined by the latent source and partially by autocorrelated independent noise.

The presynaptic firing rates were computed as follows:

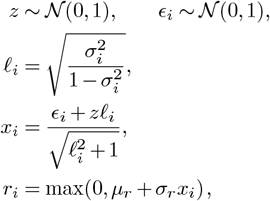

with *µ*_*r*_ = 20 Hz and *σ*_*r*_ = 10 Hz. This produces input channels with variable coupling to a shared latent source from high to low *σ*, all with equal overall variance (except for distortions due to the ReLU nonlinearity). *σ* ranged from 0 to 0.4, linearly spaced.

Presynaptic inputs were routed to the postsynaptic neuron via independent synapses (like a single presynaptic axon making multiple synapses). The proximal compartment had 1000 total synapses, such that each input was represented by 25 synapses. Distal-simple and distal-complex compartments had 40 synapses each, such that each input mapped to a single synapse. We used this structure so that the postsynaptic cell was dominated by the proximal compartment, and the distal-simple and distal-complex synapses were present to reveal how varying the strength of depression affects synaptic tuning. Inhibitory input was a non-plastic direct Poisson population (200 synapses at 10 Hz).

For each distal-complex depression ratio (*ρ*_distal_ ∈ {1.0, 1.025, 1.05, 1.075, 1.1}), we ran 10 repeats with 3 neurons per repeat for 9600s each. Proximal and distal-simple compartments were held at *ρ* = 1.1. To estimate asymptotic tuning, we averaged weights over the final 20% of each run, then normalized by both compartment-specific *w*_max_ and the number of synapses per presynaptic input.

To summarize tuning, we fit normalized weight-versus-correlation profiles with

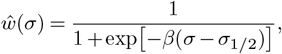

where *σ*_1*/*2_ is the sigmoid half-point used in Figure 4 as the primary summary metric.

### Visual edge tuning model, Figure 5 & 6

We modeled visual input integration of primary visual cortex by constructing a 3×3 grid of pixels, each with a random orientation. Edges appeared randomly with probability *p*(*edge*) that were centered on the central pixel. Any pixel not participating in an edge had random orientation. We sampled four evenly spaced orientations (0, *π/*4, *π/*2, 3*π/*4), so there were 36 total presynaptic input classes.

Presynaptic inputs followed von-mises tuning for orientation. Their firing rate was entirely determined by a single pixel. The firing rate was computed as:

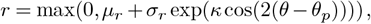

where *θ* is the orientation of a pixel, *θ*_*p*_ is the preferred orientation of the presynaptic input, and *κ* is the concentration parameter of the von-mises distribution. We set *κ* = 1.0 for all presynaptic inputs. Each input had a baseline firing rate of *µ*_*r*_ = 20 Hz and an orientation-driven firing rate of *σ*_*r*_ = 10 Hz.

The proximal compartment was only connected to the 4 inputs of the central pixel and had 200 synapses, 50 connected to each orientation. The distal-simple and distal-complex compartments were connected to all 9 pixels, each with 720 total synapses, so 20 connected to each presynaptic input.

To classify tuning in a compact way, we defined three receptive field groups (“RF Groups”). The central-preferred inputs were those in the central pixel that matched the orientation preference of the proximal synapses. The co-axial inputs were those in the boundary pixels that completed the edge of the central-preferred input (see Figure 5a or Figure 6c for illustration). All other inputs were classified together. We normalized the weight of each group to the maximum possible weight for each group to represent how strong the tuning is relative to how strong it could be in Figure 6d, e.

### Reproducibility and code availability

All code and data are available at https://github.com/landoskape/plasticity-modeling. The repository is organized and clearly documented to enable easy reproduction of results. Refer to the README.md file for instructions on how to run the code.

## ACKNOWLEDGEMENTS

We thank Jasmine Reggiani for helpful discussions and feedback on the project.

For the purpose of open access, the authors have applied a Creative Commons Attribution (CC BY) license to any Author Accepted Manuscript version arising.

## AUTHOR CONTRIBUTIONS

ATL: conceptualization, experimentation, analysis, visualization, writing

BLS: conceptualization, funding acquisition

CC: supervision, analysis, visualization

